# NS5A domain I antagonises PKR to facilitate the assembly of infectious hepatitis C virus particles

**DOI:** 10.1101/2022.08.22.504903

**Authors:** Shucheng Chen, Mark Harris

**Affiliations:** School of Molecular and Cellular Biology, Faculty of Biological Sciences and Astbury Centre for Structural Molecular Biology, University of Leeds, Leeds, LS2 9JT, United Kingdom

**Keywords:** hepatitis C virus, non-structural 5A (NS5A), genome replication, virus assembly, PKR, IRF1

## Abstract

Hepatitis C virus NS5A is a multifunctional phosphoprotein comprised of three domains (DI, DII and DIII). DI and DII have been shown to function in genome replication, whereas DIII has a role in virus assembly. We previously demonstrated that DI in genotype 2a (JFH1) also plays a role in virus assembly, exemplified by the P145A mutant which exhibited no defect in genome replication, but blocked infectious virus production (1). Here we extend this analysis to identify two other conserved and surface exposed residues proximal to P145 (C142 and E191) that shared the same phenotype. Further analysis revealed changes in the abundance of dsRNA, the size and distribution of lipid droplets (LD) and the co-localisation between NS5A and LDs in cells infected with these mutants, compared to wildtype. In parallel, to investigate the mechanism(s) underpinning this role of DI, we assessed the involvement of the interferon-induced double-stranded RNA-dependent protein kinase (PKR). In PKR-knockout cells, C142A and E191A exhibited levels of infectious virus production, LD size and co-localisation between NS5A and LD that were indistinguishable from wildtype. Co-immunoprecipitation and in vitro pulldown experiments confirmed that wildtype NS5A domain I (but not C142A or E191A) interacted with PKR. We further showed that the assembly phenotype of C142A and E191A was restored by ablation of interferon regulatory factor-1 (IRF1), a downstream effector of PKR. These data suggest a novel interaction between NS5A DI and PKR that functions to evade an antiviral pathway that blocks virus assembly through IRF1.

**Author summary:** The non-structural 5A protein (NS5A) of hepatitis C virus (HCV) plays a critical role in both virus genome replication and the assembly of infectious virus particles. NS5A is a target for potent and highly efficacious direct acting antivirals used extensively for HCV treatment. NS5A comprises 3 domains. Here, we show that the N-terminal domain I (DI) plays a role in virus assembly. Mutations in DI block both virus assembly and the perturbation of lipid droplet morphology that is a hallmark of HCV infection. Strikingly, this phenotype is abrogated by silencing of the cellular cytoplasmic double-stranded RNA sensor, PKR, a key antiviral factor. These mutations have therefore uncovered a hitherto uncharacterised antiviral pathway controlled by PKR which functions to block assembly of infectious virus particles.

## Introduction

Hepatitis C Virus (HCV) is an enveloped virus in the Flaviviridae family with a positive-sense, single-stranded RNA genome (2). HCV infection is a globally prevalent public health problem, it is the leading causative agent of chronic liver diseases including cirrhosis, hepatocellular carcinoma (HCC) and liver cancer (3, 4). As of 2015, approximately 71 million individuals worldwide (1% of the global population) were estimated to be chronically infected with HCV, resulting in more than 390,000 deaths per annum from cirrhosis and HCC (5). The recent introduction of direct-acting antivirals (DAAs), small molecule inhibitors of virus genome replication, has dramatically changed the clinical landscape. DAA treatment is able to cure the majority (∼99%) of patients rapidly and with no side-effects (6).

The HCV RNA genome contains approximately 9600 nucleotides and is comprised of a 5′-untranslated region (UTR), a single open reading frame (ORF) encoding a single polyprotein of 3000 amino acids, and a 3′-UTR (7).The polyprotein is processed co- and post-translationally by cellular and viral proteases into 10 viral proteins, namely the structural proteins: Core, E1 and E2, and p7, and the non-structural proteins: NS2, NS3, NS4A, NS4B, NS5A and NS5B (8, 9).

The subject of this study, NS5A, is a RNA-binding phosphoprotein of 49 kDa, comprised of an N-terminal amphipathic α-helix (residues 1-33) that anchors the protein to cytoplasmic membranes, a structured domain 1 (DI), and two intrinsically disordered domains (DII and DIII) linked by two low-complexity sequences (LCS-1 and -2) (10–12) (Fig 1A). X-ray crystallography demonstrated that NS5A DI is a highly structured zinc-binding domain. In addition, four different conformations of DI from genotype 1a and 1b were observed, with the same monomeric unit but exhibiting different dimeric arrangements (13–15). NS5A DI was considered to be required exclusively for genome replication (3, 12, 16), however, our previous work revealed that DI was also involved in assembly of infectious virus (1). Notably, NS5A DI has been identified as the target for one class of DAAs (eg daclatasvir and velpatasvir), as judged by the fact that mutations in DI result in resistance to these DAAs. Although the mode of action of these DAAs is not understood, they inhibit virus replication with extraordinary potency (pM EC_50_ values in culture) and have been shown to independently block both virus genome replication and virus assembly with different kinetics (17), providing further support for the role of DI in virus assembly. In contrast to DII and DIII, DI exhibits high sequence homology across all hepaciviruses (18), indicating critical and conserved functions in the virus life cycle.

**Fig 1.**
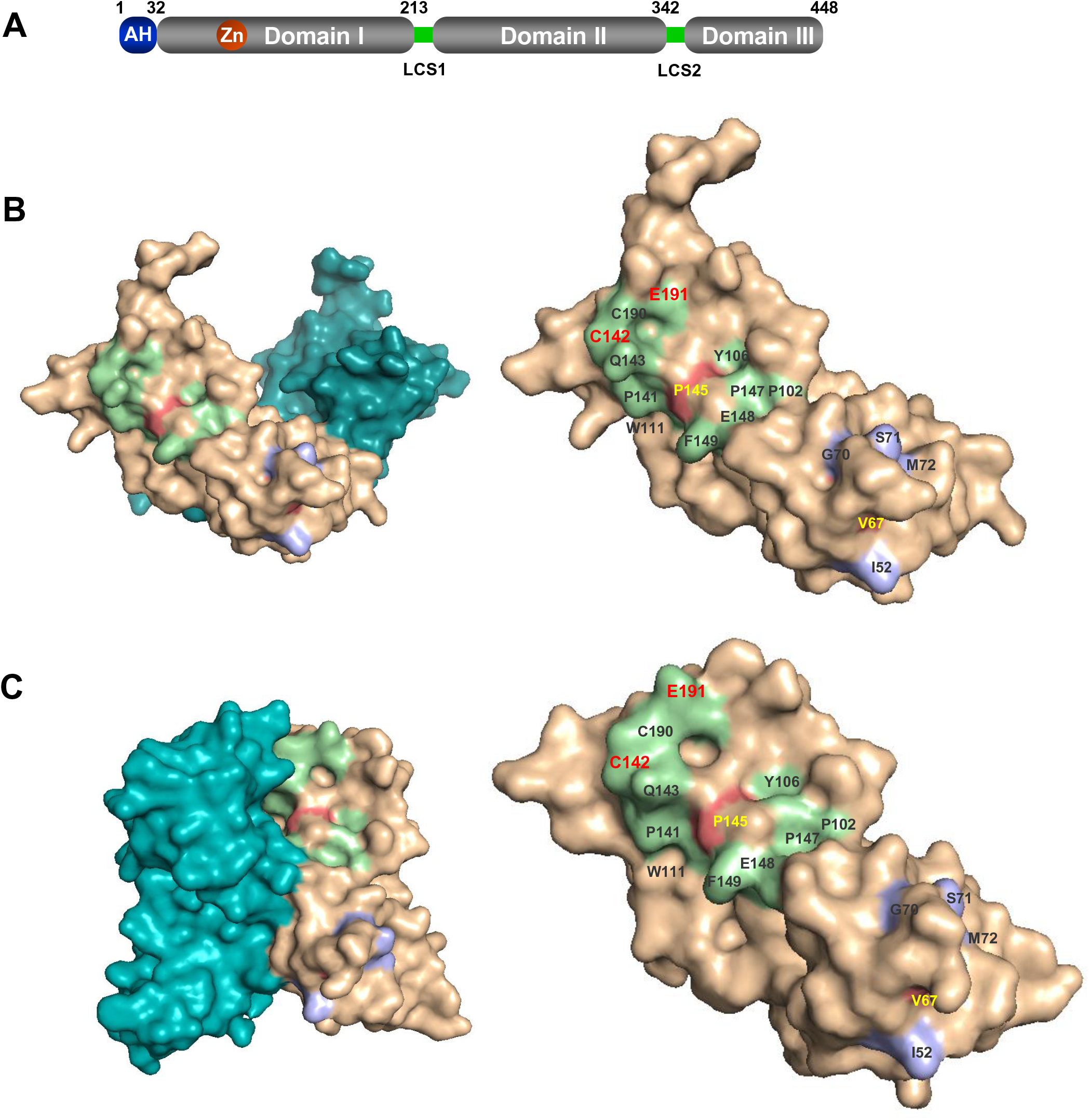
Location of mutated residues in DI. **(A)** Structure of NS5A illustrating the three domains. AH: amphipathic helix (blue), LCS: low complexity sequence (green). (**B, C)** Conserved and surface exposed residues proximal to V67 (blue) and P145 (green) are displayed in the two NS5A DI (genotype 1b) structures 1ZH1 (**B)** and 3FQM **(C)**. Dimeric forms are shown on the left, with the monomers on the right.

NS5A plays a variety of roles in the HCV life cycle. With the other non-structural proteins (NS3 to NS5B) it constitutes the viral replicase which, together with cellular factors, remodels the ER membrane to form viral replication organelles termed the membranous web (MW) (19, 20). NS5A binds directly to viral RNA (21, 22), but also modulates HCV RNA replication by interacting with other NS proteins and various cellular factors, such as vesicle-associated membrane protein-associated proteins A and B (VAP-A, VAP-B), cyclophilin A (CypA) and phosphatidylinositol-4-kinase IIIα (PI4KIIIα) (23–26).

NS5A plays an additional role in the assembly of infectious virus particles. NS5A interacts with Core (27) and both proteins are recruited to lipid droplets (LD). If this targeting is blocked either genetically or pharmacologically virus assembly is inhibited, suggesting that LDs provide a platform for virion formation (28). NS5A interacts with a number of cellular proteins that play a critical role in the assembly process. These include: diacylglycerol acyltransferase 1 (DGAT1) (29), oxysterol binding protein (OSBP) (30), phosphatidylserine-specific phospholipase A1 (PLA1A), Rab18, Tail-Interacting Protein 47 (TIP47) and Annexin A3 (31–34). However, for both genome replication and virus assembly, the precise molecular mechanisms underpinning the roles of NS5A remain elusive.

Double-stranded RNA (dsRNA)-dependent protein kinase (PKR) acts as both a sensor of virus infection by binding to viral dsRNA resulting in activation of the kinase, and also an effector via downstream consequences such as inhibition of protein translation and induction of apoptosis (35). NS5A interacts with PKR via a region in DII termed the interferon-sensitivity determining region (ISDR), the sequence of which correlates with the sensitivity of virus isolates to interferon treatment. This interaction blocks PKR activation (36, 37). The effect of PKR on the HCV lifecycle is controversial: inhibition of PKR has been shown to promote HCV genome replication and translation (38), paradoxically HCV has been reported to recruit and activate PKR to trigger induction of interferon stimulated genes (ISGs) (39, 40). Once activated, PKR has a number of downstream effects: the best characterised is the phosphorylation of the α subunit of eukaryotic initiation factor 2 (eIF2α) at Ser51 (41). This blocks translation by preventing recycling of eIF2 to the initiation complex (42). PKR also activates the transcription factor NF-κB independently of PKR catalytic activity (43, 44). PKR activation leads to dissociation of the inhibitor IκB from the p50/p65 NF-κB complex, which then enters the nucleus and activates transcription (45, 46). IFN regulatory factor 1 (IRF1) is also activated by PKR, and blocked by binding of the NS5A ISDR to PKR (47).

In this study, we confirmed the role of NS5A DI in HCV assembly. We identified two surface-exposed and conserved residues, which were not essential for virus genome replication, but disrupted the production of infectious virus particles. LD formation and the co-localisation between Core, NS5A and LDs were disrupted in cells infected with these two mutants. Intriguingly, silencing of either PKR or the downstream factor IRF1 restored the production of infectious virus, LD size and co-localisation with Core and NS5A for these two mutants. These data reveal that NS5A DI functions to allow HCV to evade inhibition of virus assembly by PKR and IRF1, and uncovers a hitherto unidentified function of PKR to inhibit virus assembly.

## Results

### Genome replication phenotypes of residues proximal to V67 and P145

NS5A DI was thought to function exclusively in HCV genome replication (12) until several years ago when we challenged this dogma and demonstrated that NS5A DI in HCV genotype 2a (JFH-1) also plays a critical role in virus assembly. This was exemplified by the phenotype of alanine substitutions at V67 and P145, which were not required for RNA replication but blocked production of infectious virus particles (1). To investigate whether the V67A/P145A phenotype was unique we identified a panel of fifteen surface-exposed and highly conserved residues proximal to V67 and P145, shown in Fig 1B/C and highlighted on the monomer structures of genotype 1b NS5A DI (PDB 1ZH1 and 3FQM) (13, 15). These residues were substituted by alanine in the context of a JFH-1 derived sub-genomic replicon (mSGR-luc-JFH-1) which contains unique restriction sites flanking the NS5A coding sequence, and a firefly luciferase reporter (48, 49). An NS5B GND mutant (polymerase-inactive) was used as a negative control. As shown in Fig 2 these mutants exhibited a number of phenotypes: Five residues proximal to V67 (I52, G70 and M72), or P145 (P141 and E148) exhibited the same phenotype as V67, ie reduced replication in Huh7 cells, which was restored to wildtype (WT) levels in Huh7.5. Six residues proximal to V67 (S71), or P145 (C142, Q143, P147, C190 and E191), were dispensable for genome replication in either Huh7 or Huh7.5 cells. In contrast, four residues proximal to P145 (P102, Y106, W111 and F149) were absolutely required for genome replication in both cell lines and were thus excluded from further analysis in this study.

**Fig 2.**
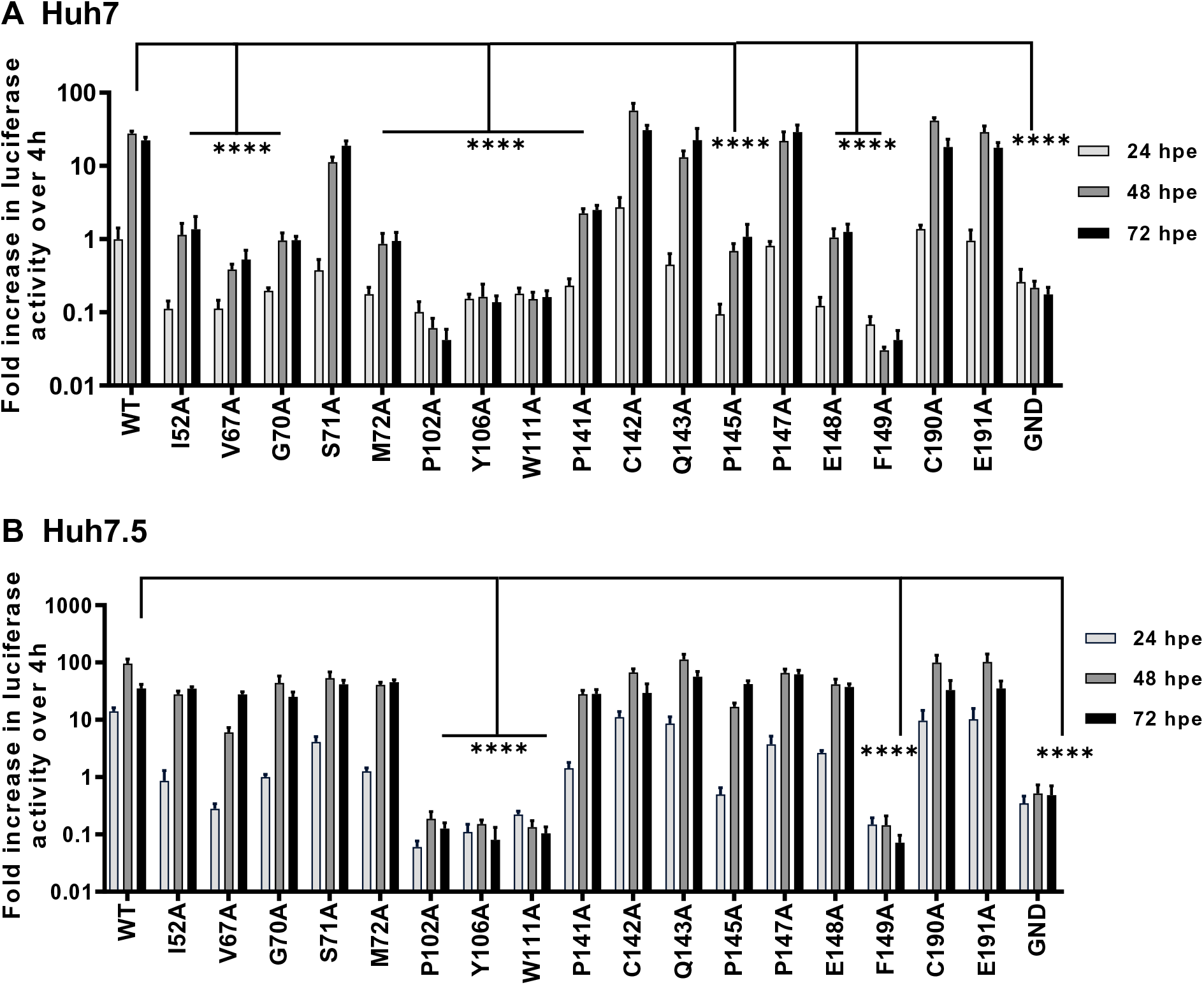
Genome replication in Huh7 and Huh7.5 cells. *In vitro* transcribed mSGR-luc-JFH-1 RNAs containing the indicated mutations were electroporated into **(A)** Huh7 and **(B)** Huh7.5 cells. Luciferase activity was measured at 4, 24, 48 and 72 hpe and the data were normalized with respect to 4 hpe.

### C142 and E191 are required for virus assembly

We next investigated whether the eleven residues that were dispensable for genome replication played a role in virus assembly and release. Alanine substitution of all of these residues were generated in the full-length mJFH-1 infectious clone (49). The assembly phenotype of these mutants was evaluated in Huh7.5 cells as in our hands they more efficiently supported both virus genome replication (Fig 2 – compare WT values in A and B), and assembly (data not shown). As shown in Supp. Fig. S1, nine of these mutants produced similar levels of released infectious virus to WT, however two of the mutants (C142A and E191A) exhibited a significant defect in virus production. We therefore focused our further investigations on these two mutants. Interestingly, C142 was shown to be connected to C190 by a disulphide bond in the ‘open’ dimer conformation (1ZH1), but not the ‘closed’ dimer (3FQM) of NS5A DI (Supp. Fig. S2). Previous mutagenesis had shown that this disulphide bond is not required for genome replication (12), consistent with our replication data (Fig 2). As C190 is positioned between C142 and E191 on the NS5A DI surface (Supp. Fig. S2), we included C190A as a control in the detailed analysis of the C142A and E191A phenotypes, as described henceforth.

We first confirmed virus genome replication for the mutant infectious clones C142A, C190A and E191A both directly by qPT-PCR (Fig 3A), and indirectly by quantifying NS5A positive cells using an IncuCyte S3 cell imager (50) (Fig 3B). Reassuringly, replication of all three mutants was indistinguishable from WT, mirroring the SGR data. Western blot analysis confirmed that NS5A and the structural proteins Core and E2 were expressed at equivalent levels to WT (Fig 3C).

**Fig 3.**
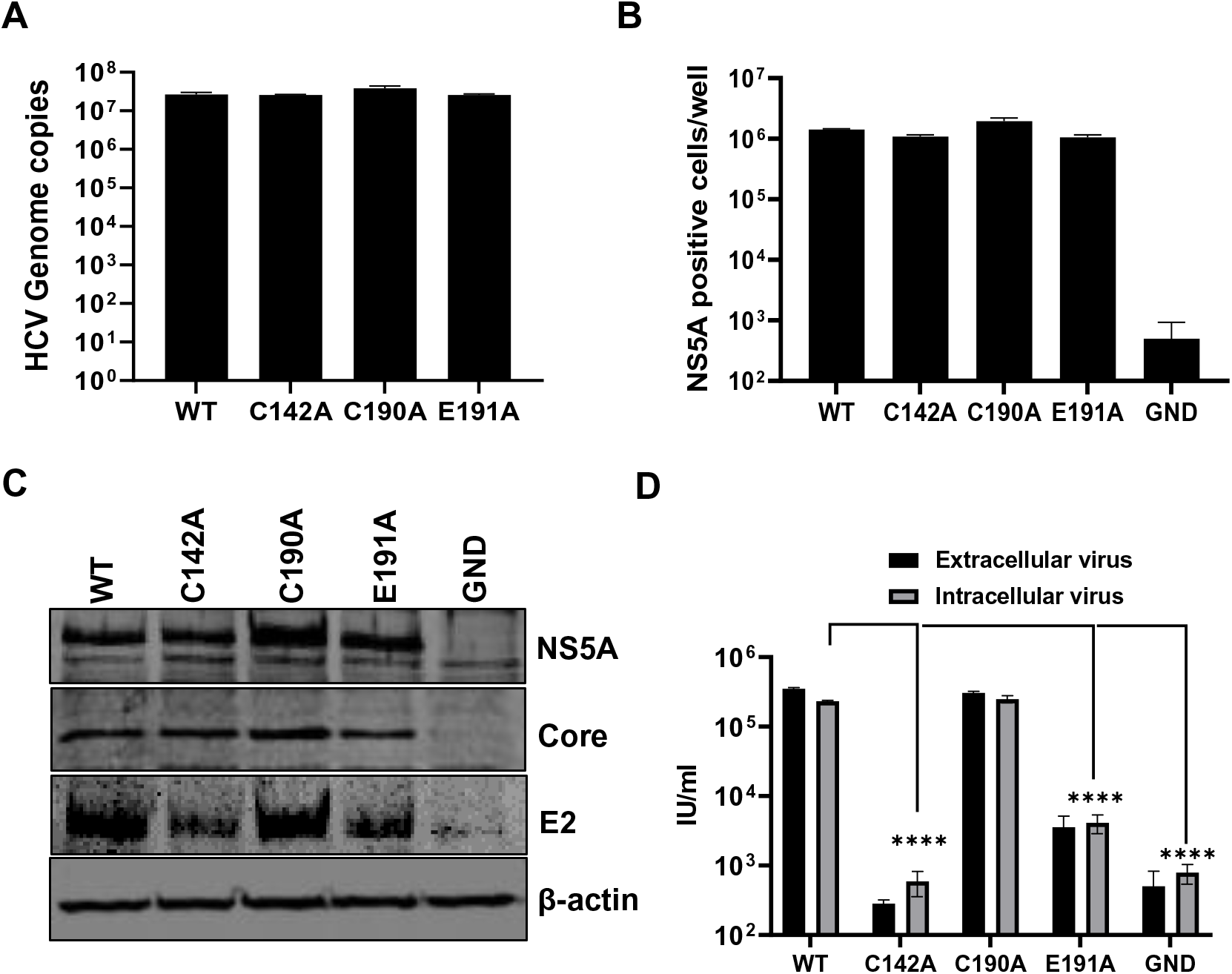
Virus assembly phenotypes in Huh7.5 cells. Huh7.5 cells were electroporated with mJFH-1 WT and DI mutant C142A, C190A and E191A RNAs, together with an NS5B GND mutant as negative control. Virus genome replication was analysed directly by quantification of genome copies in cell lysates using qRT-PCR (**A**), and indirectly by enumerating NS5A positive cells at 72 hpe using the IncuCyte S3 (**B**). (**C**) Cell lysates were collected at 72 hpe and analysed by western blotting with the indicated antibodies. (**D)**. Extra- and intracellular virus harvested at 72 hpe were titrated in Huh7.5 cells and quantified using the IncuCyte S3.

To assess both virus assembly and release we proceeded to determine intracellular and extracellular virus titres (Fig 3D). This analysis revealed that C142 was absolutely required for virus assembly with levels of both intracellular and extracellular virus indistinguishable from the negative control (NS5B GND). In contrast C190A had no effect on WT levels of infectivity, and E191A exhibited an intermediate phenotype with an approximately 2-log reduction compared to WT. We conclude that C142 and E191 play a role in virus assembly, and the fact that C190 is dispensable further suggests that the disulphide bond observed in the ‘open’ structure of DI is not required for the function of NS5A during genome replication or assembly.

### NS5A DI mutants alter the morphology and distribution of lipid droplets

To better characterise the role of C142 and E191 on infectious virus production, we used high resolution confocal microscopy (Airyscan) to observe the co-localisation between viral proteins and cellular factors. Key organelles during virus assembly are lipid droplets (LDs), to which both Core (51) and NS5A (52) are recruited. The disruption of LDs either pharmacologically or genetically (53) inhibits virus assembly. We previously showed that in cells infected with the P145A mutant virus, LDs were more abundant and smaller in size compared to WT infected cells (1). As shown in Fig 4, this phenotype was recapitulated for both C142A and E191A: cells infected with WT and C190A exhibited an average of 100 LD with a cross-sectional area of approximately 1.0 μm^2^, whereas C142A and E191A displayed >200 LD with a significantly smaller area (0.2 μm^2^), similar to mock-infected cells (Fig 5A).

**Fig 4.**
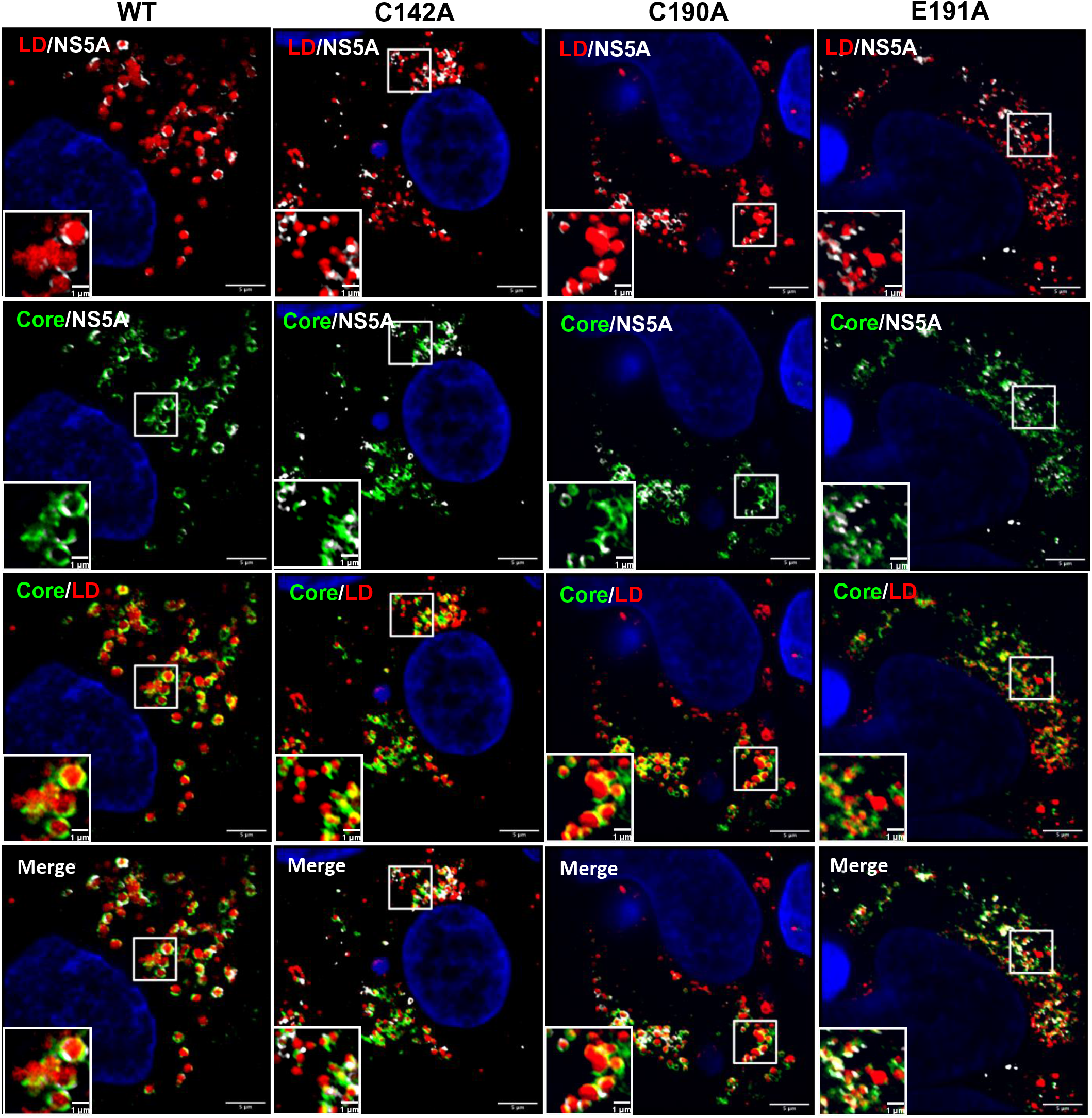
Co-localisation between NS5A, Core and LD. Huh7.5 cells were electroporated with mJFH-1 WT and DI mutant C142A, C190A and E191A RNAs and seeded on to coverslips. At 72 hpe cells were stained with sheep anti-NS5A (white), rabbit anti-Core (green), BODIPY 558/568-C12 (red) and DAPI. Co-localisation was observed using Airyscan microscopy. Representative images are presented. A representative image of mock electroporated cells is shown in Supp. Fig. S6A. Scale bars are 5 μm and 1 μm (insets).

**Fig 5.**
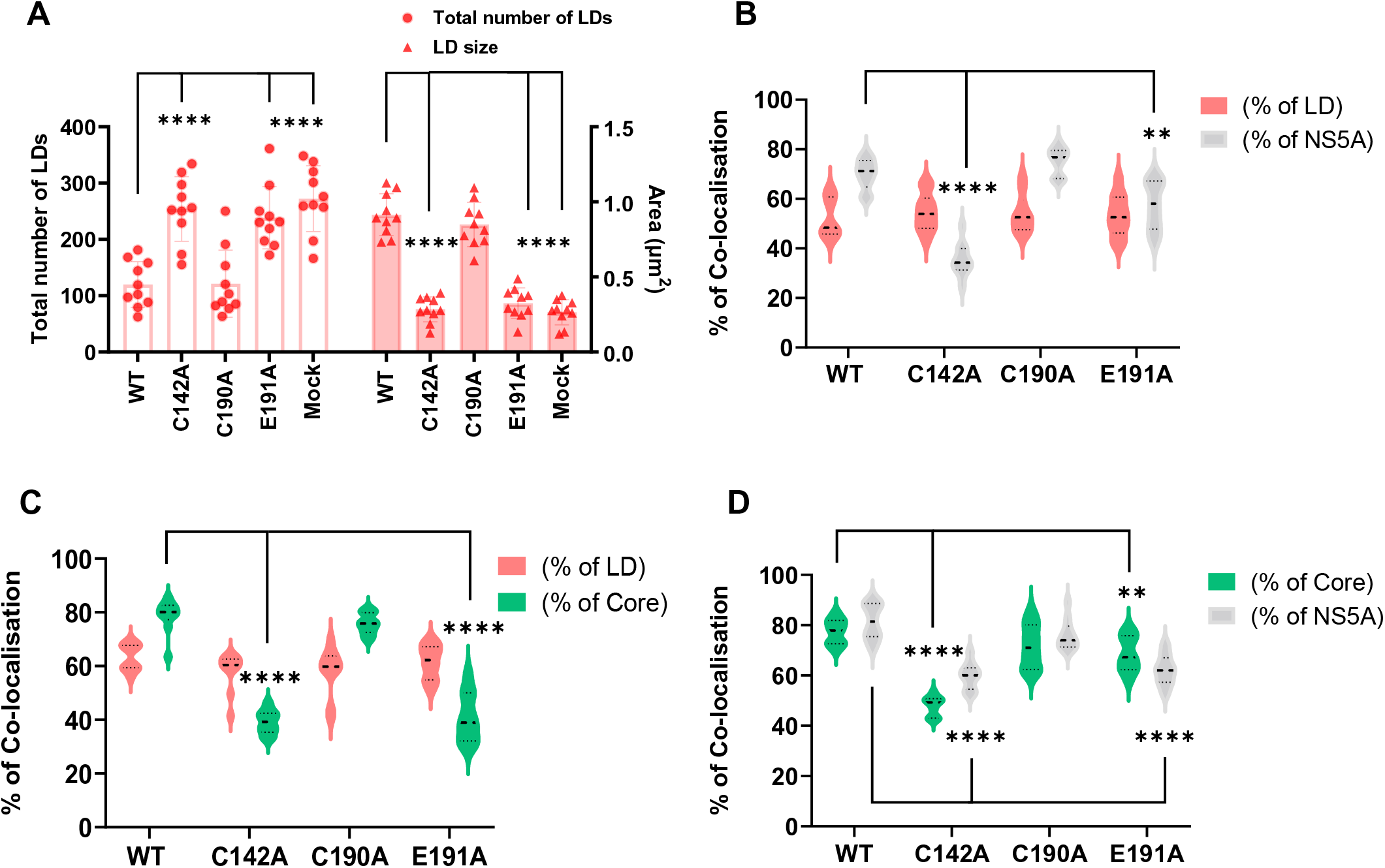
Quantification of LD size and co-localisation with NS5A and Core. **(A)** LD numbers and size in cells from Fig 4 were calculated using Analyze Particles module of Fiji. Mock: uninfected Huh7.5 cells. (**B)** Quantification of the percentage of LD colocalized with NS5A (red), and NS5A colocalized with LD (grey). (**C)** Quantification of the percentage of LDs colocalized with Core (red), and Core colocalized with LD (green). (**D**) Quantification of the percentage of Core colocalized with NS5A (green), and NS5A colocalized with Core (grey). Co-localisation was analyzed in 10 cells for each construct using Fiji. Significant differences from WT denoted by ** (P<0.01) and **** (P<0.0001).

The differences in both LD size and quantity were consistent with the assembly defective phenotypes of C142A and E191A. To extend this analysis we quantified the colocalisation of Core and NS5A with LDs. As shown in Fig 5B, the percentage of NS5A co-localising with LDs was significantly decreased for C142A and E191A compared to WT. However, the inverse value (percentage of LD colocalised with NS5A) was comparable for all mutants and WT, consistent with the suggestion that the interaction of NS5A with LD was disrupted by C142A and E191A mutations. Similar results were observed for the interaction of LD with Core (Fig 5C). Finally, we quantified the colocalisation of Core and NS5A and observed a reduction for C142A and E191A (Fig 5D). This reduction was less pronounced for E191A, consistent with the less marked phenotype of this mutant. We also assessed the distribution of LDs by quantifying their distance from the nucleus. As shown in the representative images in Fig. 6A, and quantified in Fig 6B, LDs in C142A and E191A infected cells were significantly closer to the nucleus than WT or C190A, although still more dispersed than mock infected cells. Together, these data are consistent with the proposition that NS5A DI is involved in the targeting of NS5A to LDs and functions to perturb LD morphology and distribution to promote virus assembly.

**Fig 6.**
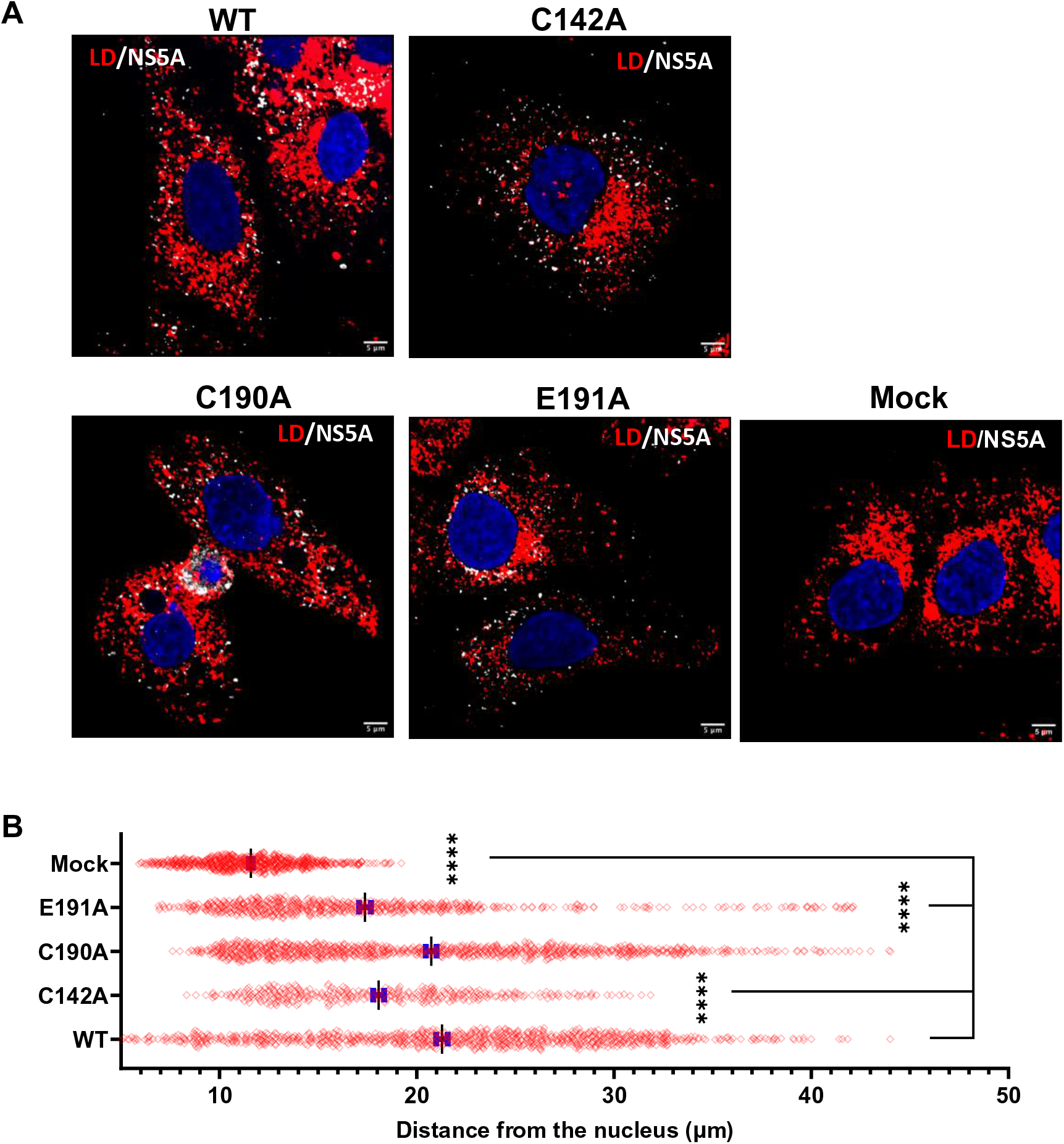
Analysis of LD distribution. **(A)** Cells were stained at 72 hpe with sheep anti-NS5A (white), BODIPY 558/568-C12 (red) and DAPI. (**B)** Distances of LDs from the nuclear membrane were evaluated using the Analyze Particles module of Fiji. Significant differences from WT denoted by **** (P<0.0001).

### PKR silencing or inhibition recovers the virus assembly phenotype of C142A and E191A

It was recently shown that CypA is critical for HCV genome replication in Huh7 cells but not Huh7.5 cells (54). Further analysis led to the conclusion that NS5A interacts with CypA to evade PKR-dependent antiviral responses. The differential sensitivity of HCV to CypA in Huh7 compared to Huh7.5 cells was reminiscent of our initial observation that P145A fails to undergo genome replication in Huh7 cells, but is only modestly impaired in Huh7.5 cells (1) (Fig 2). This led us to assess whether either CypA and/or PKR play a role in virus assembly and whether the NS5A DI mutants studied thus far can shed light on such mechanisms. Consistent with previous studies, we first confirmed that silencing of CypA or PKR in Huh7.5 cells had no effect on genome replication (Supp. Fig. S3A). We did however, note that the production of infectious virus was unaffected by PKR silencing but reduced by ∼100-fold in CypA shRNA cells (Supp. Fig. S3B).

We proceeded to analyse the genome replication and assembly of the 3 DI mutants in PKR silenced Huh7.5 cells. As expected, genome replication (Fig 7B) and viral protein production (Fig 7C, D) were unaffected by the lack of PKR. Indeed, overall levels of genome replication as measured by qRT-PCR modestly increased compared to Huh7.5 cells (compare Fig 7B to Fig 3A). A surprising picture emerged when we analysed the assembly and release of the mutants: production of infectious virus by C142A and E191A was restored to the same level as WT and C190A in PKR silenced Huh7.5 cells (Fig 7E). To further confirm that this was due to the lack of PKR, as opposed to an off-target effect of the sgRNA, we treated Huh7.5 cells with the small molecule PKR inhibitor C16 at 24 h post electroporation. Reassuringly, pharmacological inhibition of PKR function also restored the production of infectious virus by C142A and E191A (Fig 7F). These data are consistent with a role for PKR in blocking virus assembly and point to a role of NS5A DI in antagonising this previously undefined function of PKR.

**Fig 7.**
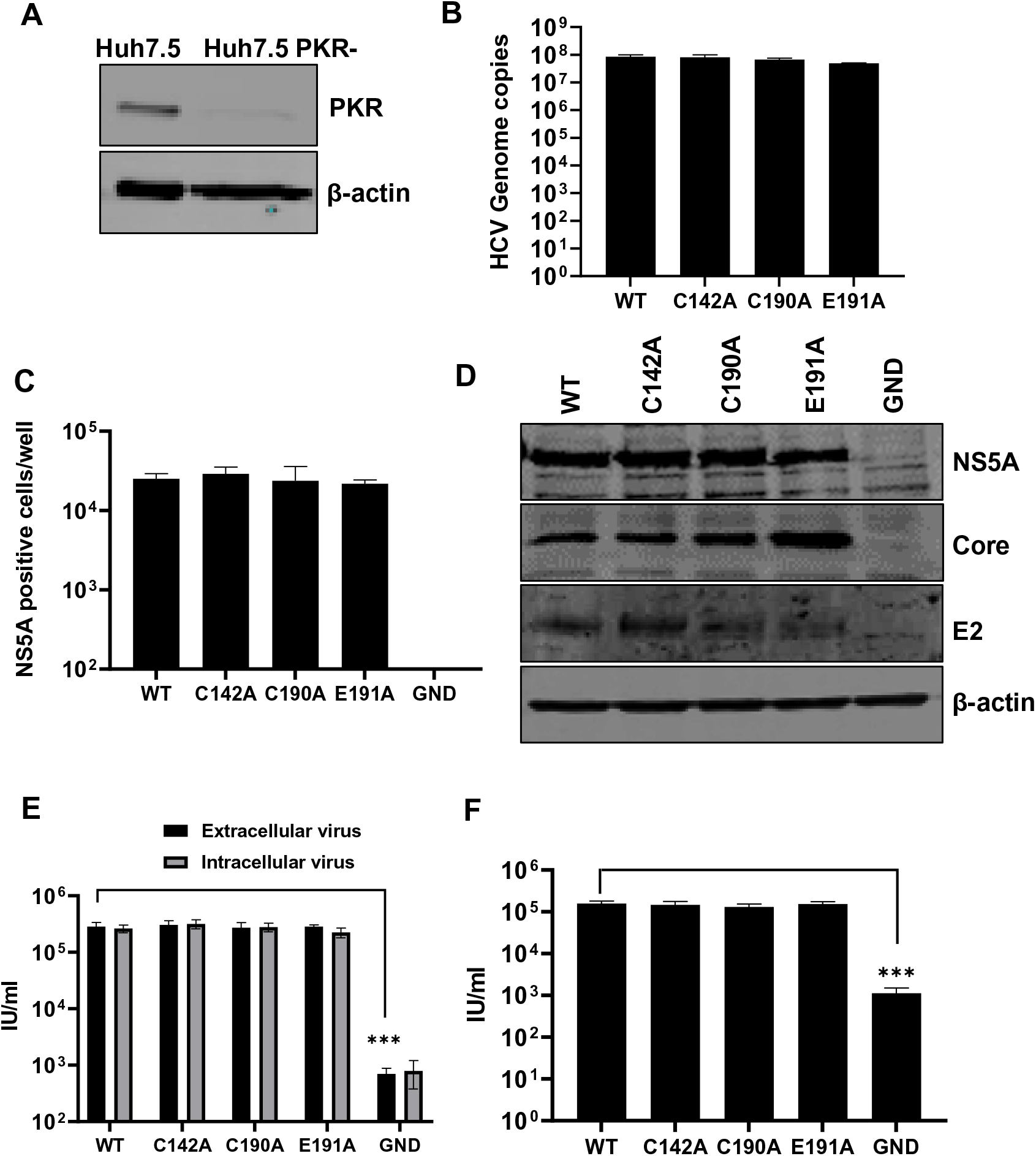
Virus assembly in PKR-silenced Huh7.5 cells. **(A)** PKR expression was detected in silenced Huh7.5 cells by western blot. Huh7.5 cells were electroporated with mJFH-1 WT and DI mutant C142A, C190A and E191A RNAs, together with an NS5B GND mutant as negative control. Virus genome replication was analysed directly by quantification of genome copies in cell lysates using qRT-PCR (**B**), and indirectly by enumerating NS5A positive cells at 72 hpe using the IncuCyte S3 (**C**). **(D)** Cell lysates were collected at 72 hpe and analysed by western blotting with the indicated antibodies. **(E)** Extra- and intracellular virus harvested at 72 hpe were titrated in Huh7.5 cells and quantified using the IncuCyte S3. (**F)** Electroporated cells were treated with the PKR inhibitor C16 from 24 hpe, extracellular virus was harvested at 72 hpe and titrated as described in **(E).**

### PKR silencing restores the LD phenotype of the assembly mutants

We next sought to determine whether the restoration of infectious virus production by PKR silencing was associated with concomitant changes in LD morphology, distribution and the association with NS5A and Core. We therefore repeated the confocal analysis as described in Figs 4-6 in PKR silenced cells. As shown in the representative images in Fig 8, and quantified in Fig 9, no differences between WT and the three mutants with regard to LD number and size were observed (Fig 9A), although it should be noted that overall the size of LDs in infected cells was slightly reduced (Fig 5A). Co-localisation analysis also revealed that, unlike in Huh7.5 cells, no differences were observed between WT or the three mutants in terms of their co-localisation between NS5A and LD (Fig 9B), Core and LD (Fig 9C) or NS5A and Core (Fig 9D).

**Fig 8.**
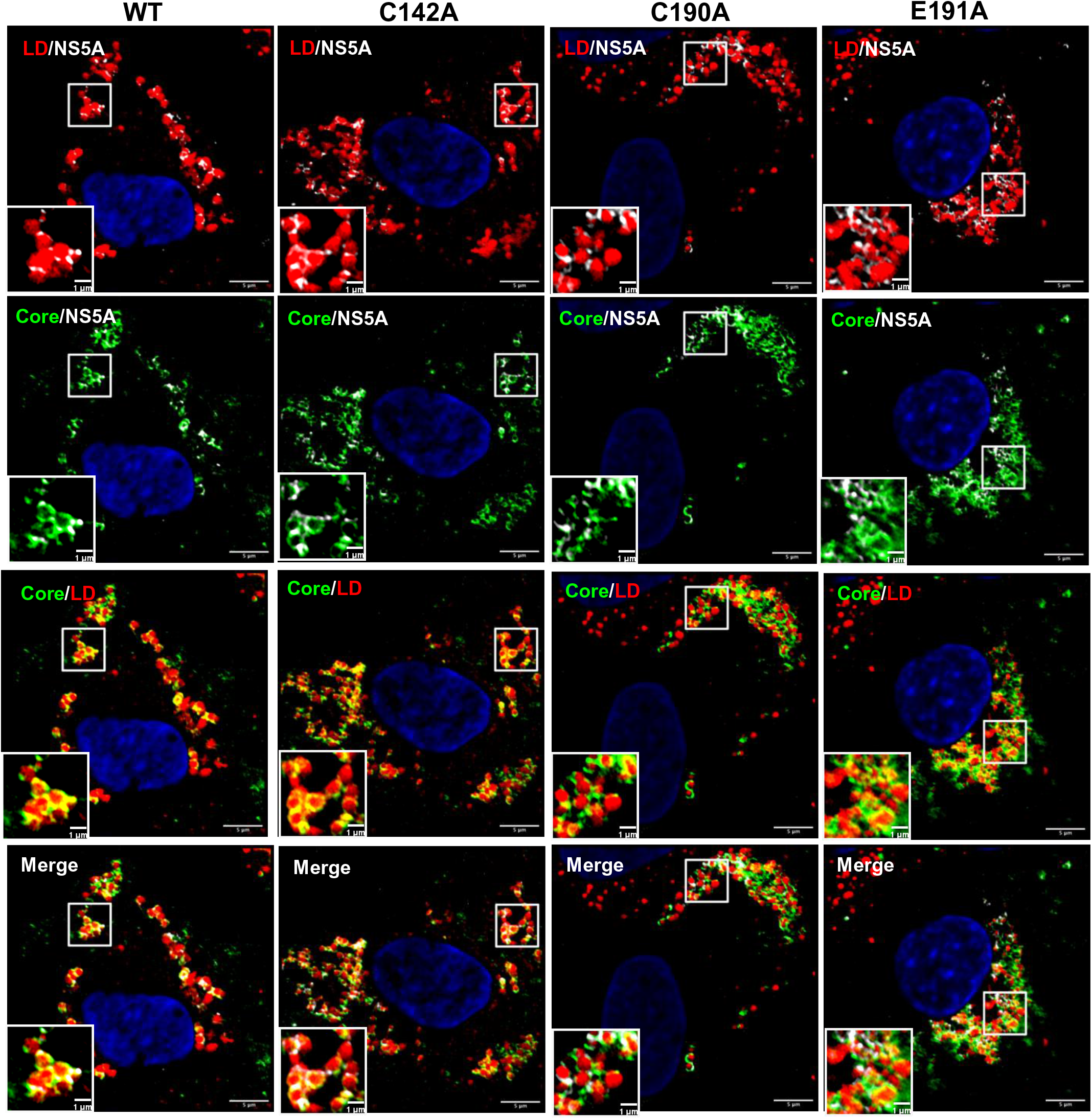
Co-localisation between NS5A, Core and LD in PKR-silenced Huh7.5 cells. Huh7.5 cells silenced for PKR expression were electroporated with mJFH-1 WT and DI mutant C142A, C190A and E191A RNAs and seeded on to coverslips. Cells were stained at 72 hpe with sheep anti-NS5A (white), rabbit anti-Core (green), BODIPY 558/568-C12 (red) and DAPI. Co-localisation was observed using Airyscan microscopy. Representative images are shown. Representative images of mock electroporated cells are shown in Supp. Fig. S6B. Scale bars are 5 μm and 1 μm (insets).

**Fig 9.**
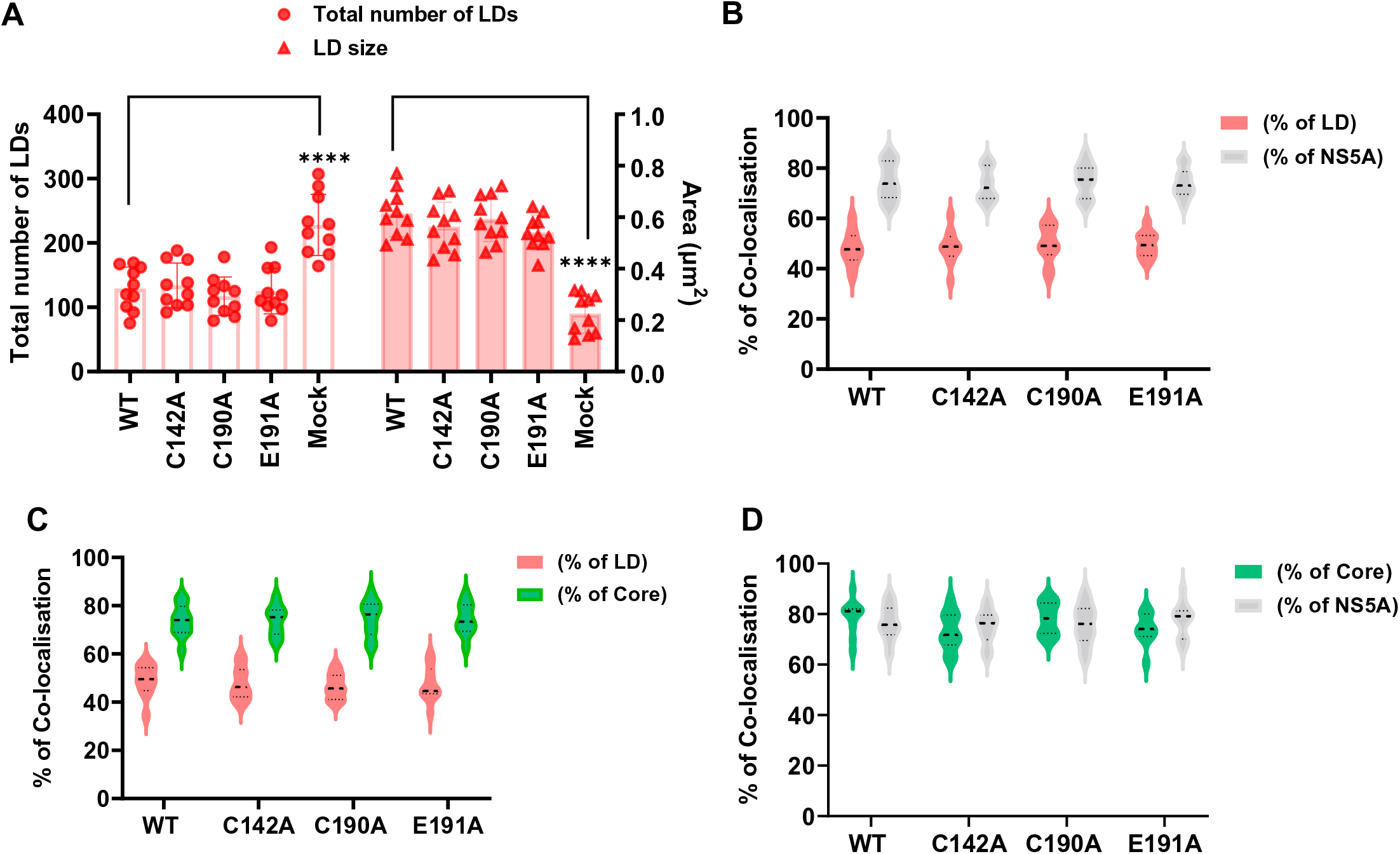
Quantification of LD size and co-localisation with NS5A and Core in PKR-silenced Huh7.5 cells. **(A)** LD numbers and size from Fig 8 were calculated using Analyze Particles module of Fiji. Mock: uninfected Huh7.5 cells. (**B)** Quantification of percentage of LDs colocalizing with NS5A (red), and NS5A colocalized with LD (grey). (**C)** Quantification of the percentage of LD colocalized with Core (red), and Core colocalized with LD (green). (**D**) Quantification of the percentage of Core colocalized with NS5A (green), and NS5A colocalized with Core (grey). Co-localisation was analyzed in 10 cells for each construct using Fiji. Significant differences from WT denoted by ns (P>0.05) and **** (P<0.0001).

Lastly, we assessed the distribution of LDs by quantifying their distance from the nucleus. As shown in the representative images in Fig. 10A, and quantified in Fig 10B, the distribution of LDs in PKR silenced cells was comparable to that observed in Huh7.5 cells: in C142A and E191A infected cells LDs were significantly closer to the nucleus than WT and C190A, although still more dispersed than mock infected cells. These data are consistent with the suggestion that PKR functions to disrupt virus assembly by blocking the HCV-mediated perturbations of LD morphology. Furthermore, NS5A DI antagonises this function of PKR through the surface exposed residues C142 and E191.

**Fig 10.**
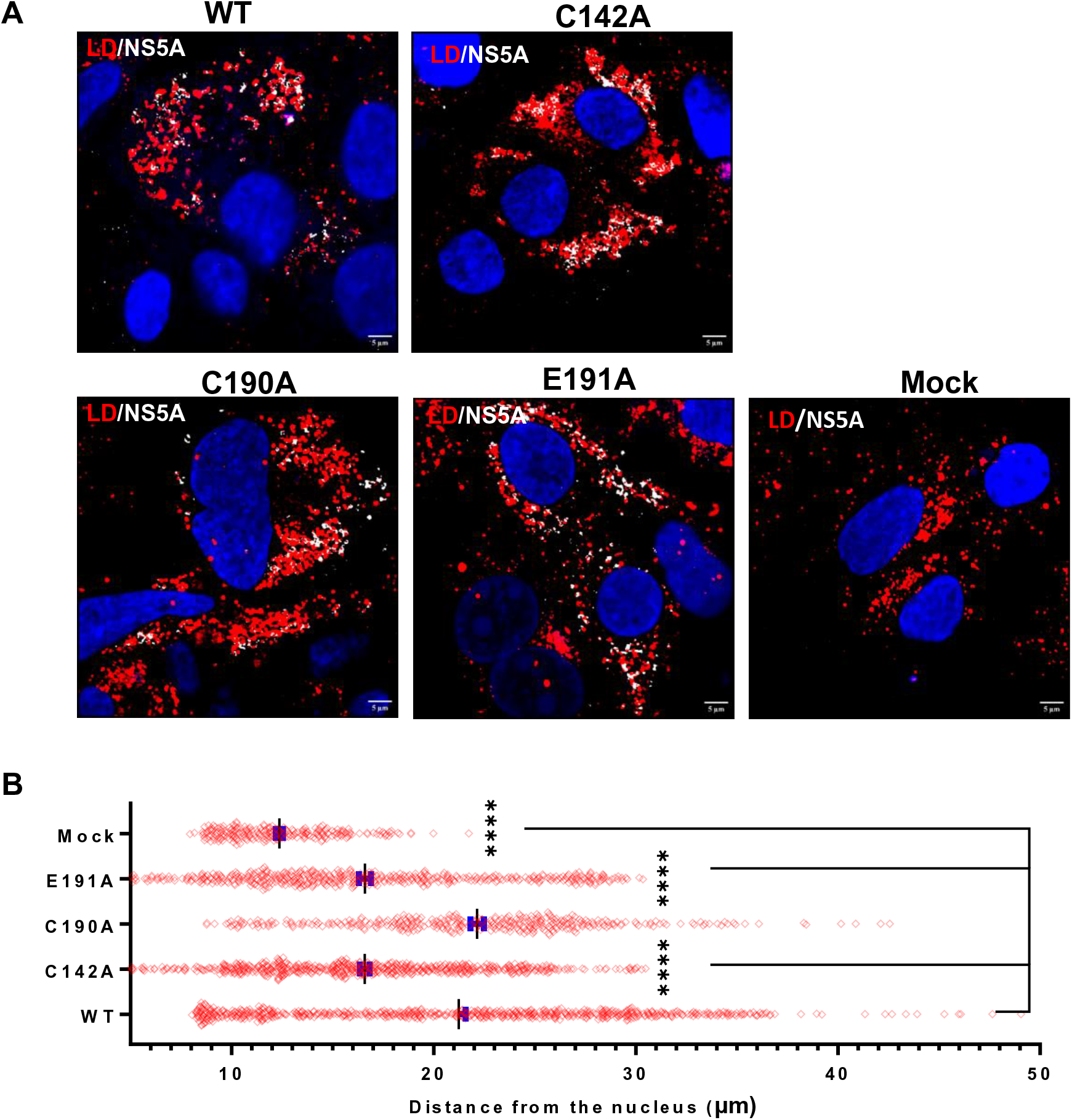
Analysis of LD distribution in PKR-silenced Huh7.5 cells. **(A)** Cells were stained at 72 hpe with sheep anti-NS5A (white), BODIPY 558/568-C12 (red) and DAPI. (**B)** Distance of LDs from the nucleus was evaluated using the Analyze Particles module of Fiji. Significant differences from WT denoted by **** (P<0.0001).

### Assembly defective mutants C142A and E191A exhibit reduced dsRNA abundance

We next sought to understand the differences between WT and the NS5A DI assembly-defective mutants with regard to PKR activation. PKR is activated by binding to dsRNA via two N-terminal RNA-binding domains (dsRBD) (55, 56). This leads to dimerisation of PKR, autophosphorylation and activation of the C-terminal kinase domain. dsRNA is generated as a replication intermediate and co-localises with NS5A, Core and LDs (57). Furthermore, the HCV genome contains many structured RNA elements with extensive double-stranded regions (58). Although genome replication is likely protected from PKR as it occurs within the membranous web (19), nascent genomes must be transported through the cytoplasm to sites of assembly and during this process may be detected by PKR. We therefore assessed the co-localisation between dsRNA, NS5A and LDs in Huh7.5 cells using a well-characterised dsRNA-specific antibody, J2 (57). As expected, in WT infected cells, we observed co-localisation of dsRNA with both NS5A and LDs (Fig 11). This co-localisation was also quantified in cells infected with the three mutants and surprisingly, revealed no significant differences in the co-localisation of NS5A and dsRNA (Fig 12A) or LD and dsRNA (Fig 12B). However, the number of dsRNA foci in C142A and E191A were reduced compared to WT and C190A (Fig 12C-D). WT and C190A infected cells exhibited 200-300 dsRNA punctae whereas C142A and E191A infected cells had between 100-200 punctae. Given that the overall levels of HCV genomes in all of these cells were equivalent (Fig 3A), this suggests that the majority of dsRNA foci actually represent nascent genomes, rather than replicative intermediates. For the two assembly defective mutants, C142A and E191A, we propose that our results are consistent with the failure of NS5A to block PKR binding to dsRNA elements in nascent genomes, resulting in PKR activation, dsRNA degradation and stimulation of an antiviral response.

**Fig 11.**
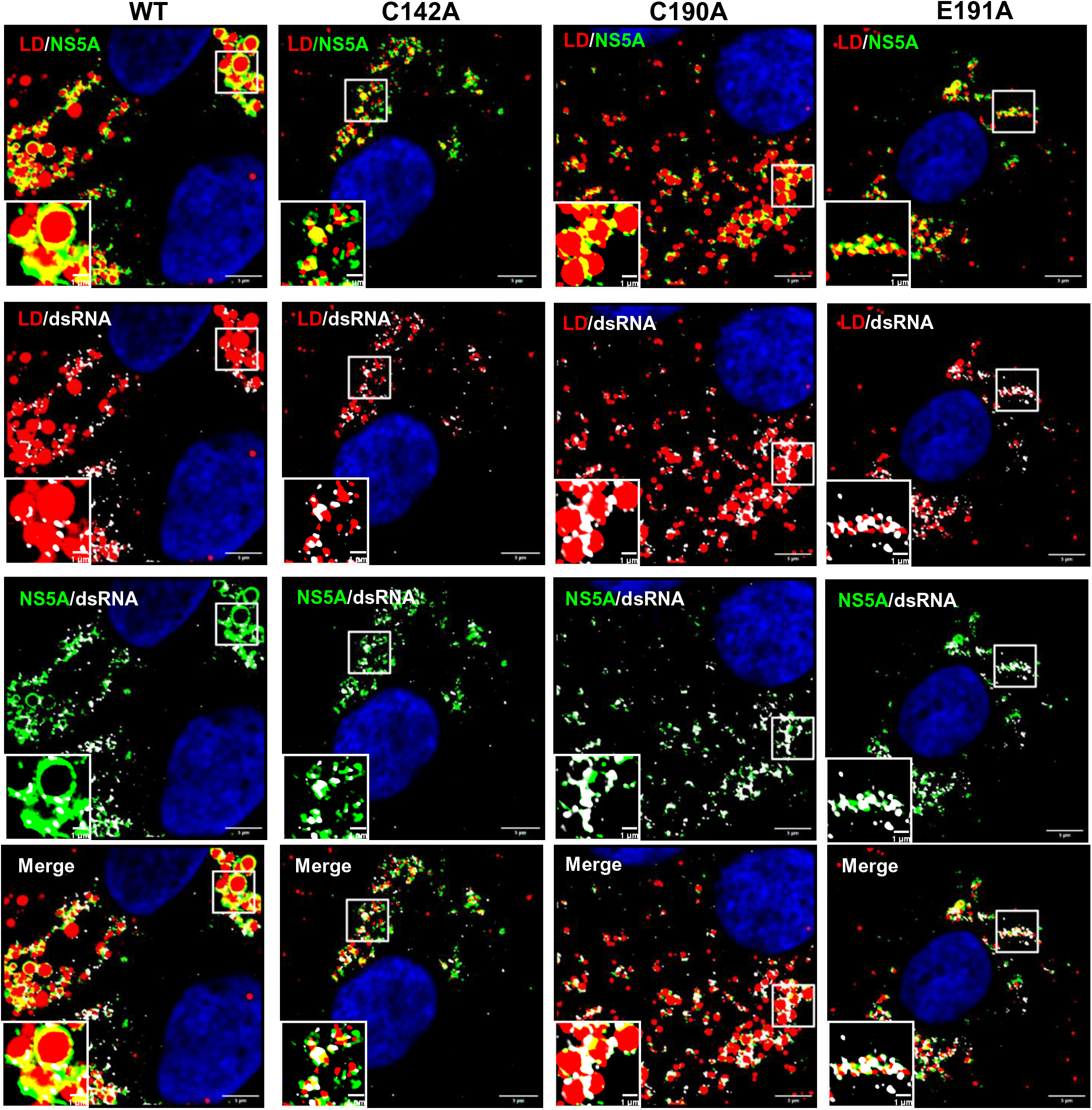
Co-localisation of NS5A, dsRNA and LD. Huh7.5 cells were electroporated with mJFH-1 WT and DI mutant C142A, C190A and E191A RNAs and seeded on to coverslips. At 72 hpe cells were stained with sheep anti-NS5A (green), mouse anti-dsRNA J2 (white), BODIPY 558/568-C12 (red) and DAPI. Co-localisation was observed using Airyscan microscopy. Representative images are shown. Representative images of mock electroporated cells is shown in Supp. Fig. S6C. Scale bars are 5 μm and 1 μm (insets).

**Fig 12.**
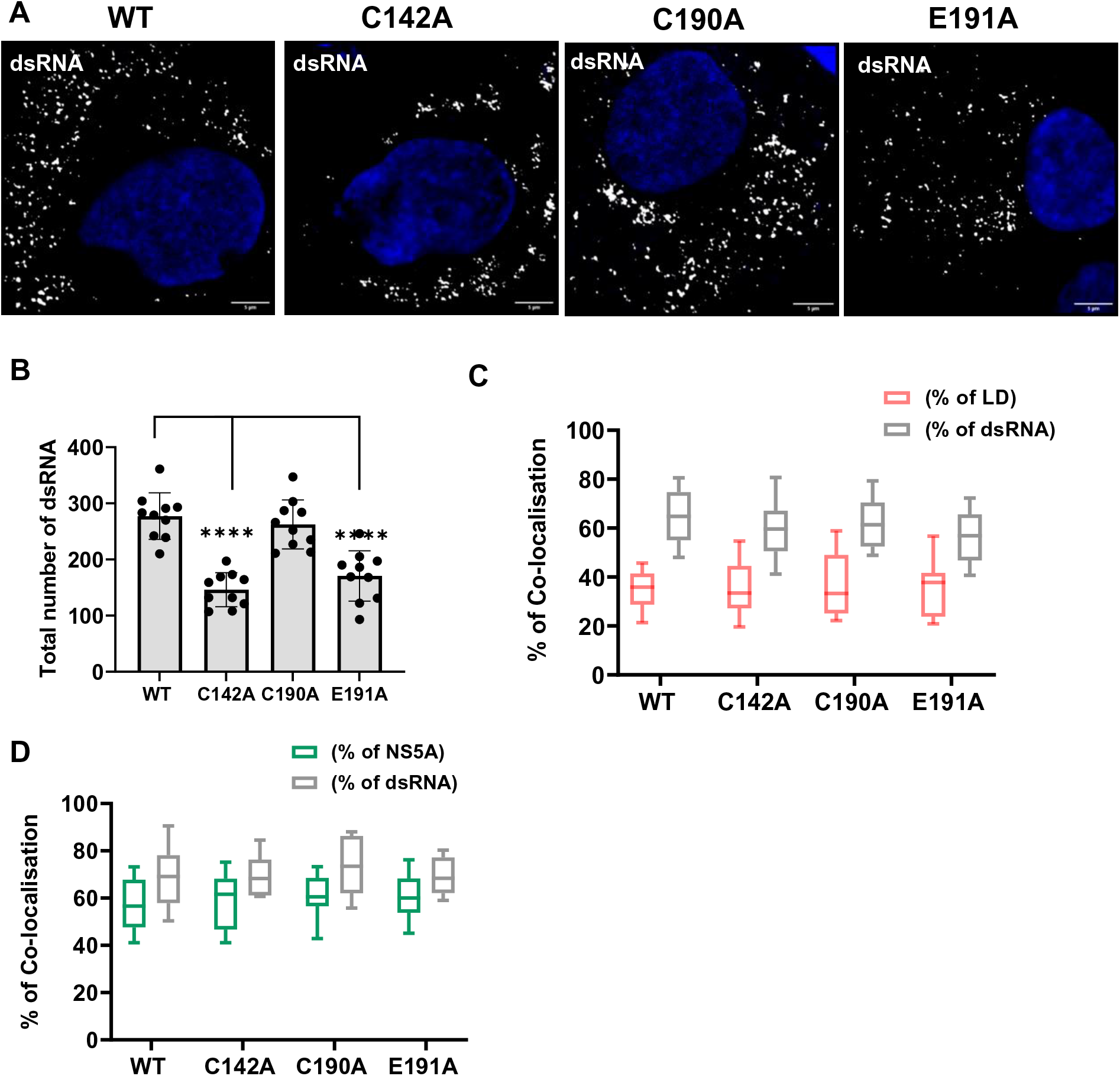
Quantification of dsRNA punctae and co-localisation with NS5A and LD. (**A)** Quantification of the percentage of LD colocalized with dsRNA (red), and dsRNA colocalized with LD (grey). (**B)** Quantification of the percentage of NS5A colocalized with dsRNA (green), and dsRNA colocalized with NS5A (grey). Co-localisation was analyzed in 10 cells from Fig 11 using Fiji. **(C)** Representative images of Huh7.5 cells electroporated with mJFH-1 WT and DI mutants C142A, C190A and E191, seeded on to coverslips and stained at 72 hpe with mouse anti-dsRNA J2 (white) and DAPI. Scale bars are 5 μm. **(D)**. Numbers of dsRNA punctae of each sample from **(C)** were calculated using the Analyze Particles module of Fiji.

### NS5A DI interacts with PKR

NS5A has been previously demonstrated to directly interact with PKR via a region in D2 termed the interferon sensitivity determining region (ISDR) (37). These studies were performed with NS5A from genotype 1b and showed exquisite sensitivity to the amino acid sequence of the ISDR. The homology in this region between genotype 1b and JFH-1 (genotype 2a) is low (44% identity) so it is unlikely that the ISDR in JFH-1 binds to PKR, although this has not been formally proven. We considered that the ability of NS5A DI to block PKR could be explained by a direct interaction between the two proteins, dependent on C142 and E191. To test this we immunoprecipitated PKR from Huh7.5 cells electroporated with either WT or the three mutants and investigated the presence of NS5A in the immunoprecipitates by western blotting. As shown in Fig 13A, only WT and C190A NS5A co-precipitated with PKR, whereas C142A and E191A NS5A did not. We also investigated whether NS5A was able to bind to activated PKR by performing immunoprecipitations with an antibody to phosphorylated PKR (P-PKR). Although overall levels of P-PKR were low, this analysis clearly showed that (as for the total PKR) only WT and C190A NS5A co-precipitated with P-PKR (Fig 13B).

**Fig 13.**
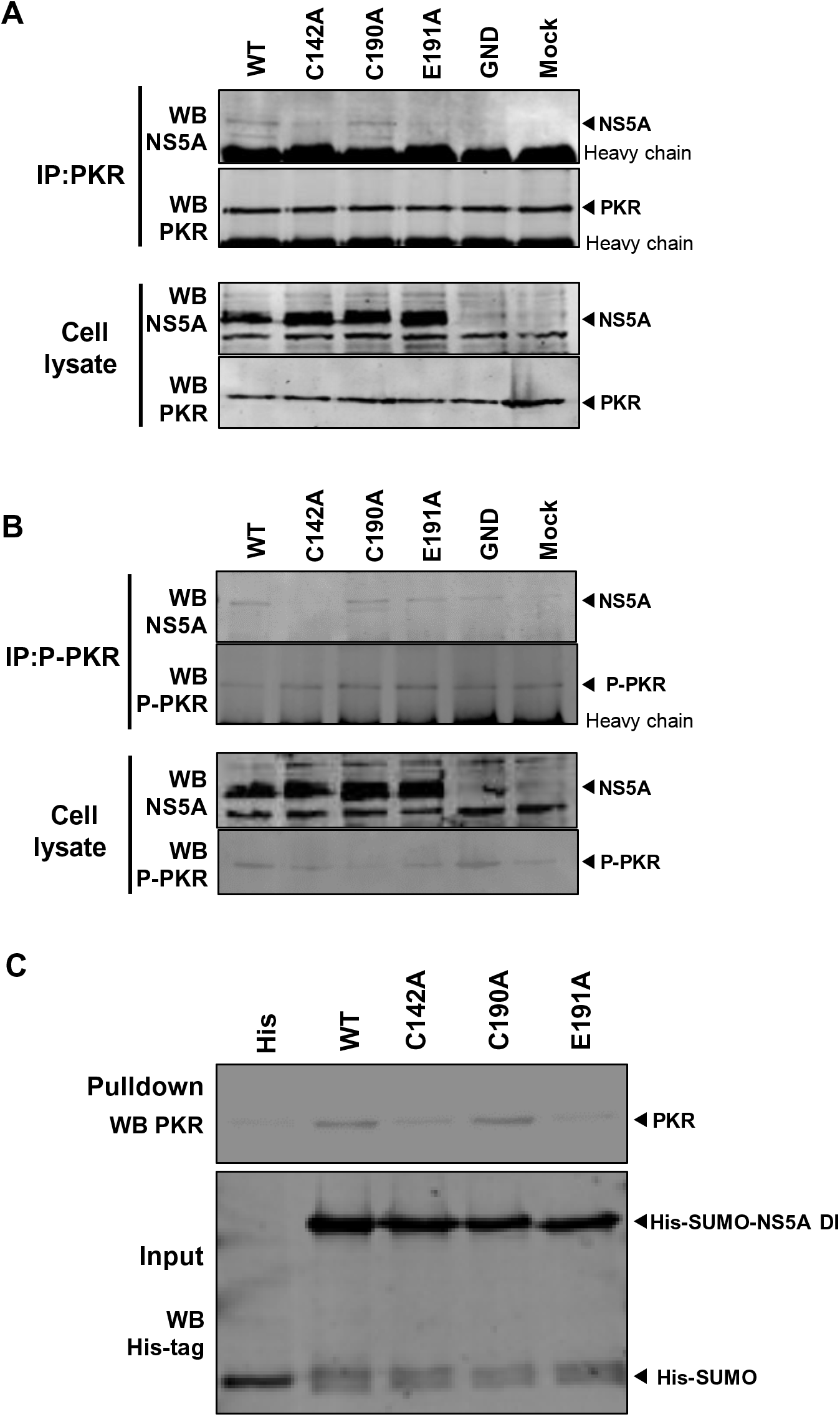
Protein-protein interaction analysis of NS5A and PKR. **(A, B)** Huh7.5 cells electroporated with mJFH-1 WT and DI mutants C142A, C190A and E191 were lysed and immunoprecipated with anti PKR antibody (**A**) or anti phospho-PKR antibody (**B**). Immunoprecipitates (top) and lysates (bottom) were analysed by western blot with the indicated antibodies. (**C)** His-SUMO tagged NS5A DI and mutants were purified and bound to Dynabead His-Tag beads as a bait to precipitate PKR protein which was overexpressed in HEK293T cells. Pulldowns were analysed by western blotting for PKR (top), and inputs verified by western blotting for the His-tag (bottom).

To further validate the interaction between NS5A DI and PKR, His-SUMO tagged NS5A DI (WT and mutants) were expressed in *E.coli*, purified and used as bait to precipitate PKR which was overexpressed in HEK293T cells. Confirming the co-immunoprecipitation data, only WT and C190A DI, but not C142A and E191A, were able to precipitate PKR from the cell lysate (Fig 13C), confirming that DI is indeed able to bind to PKR. These data are consistent with the hypothesis that NS5A DI binds directly to PKR to prevent it activating downstream pathways that would lead to an antiviral response against virus assembly. We therefore turned our attention to identifying the downstream PKR effector(s) responsible.

### A role for the PKR effector IRF1 in blocking HCV assembly

A well characterised downstream effector of PKR is the phosphorylation of eIF2α at Ser51 to block protein synthesis. PKR also activates NFκB independently of its catalytic activity (43, 44), as well as interferon regulatory factor 1 (IRF1). Although the mechanism by which PKR activates IRF1 is uncharacterised, unlike NFκB activation it is dependent on PKR catalytic activity (59).

Western blotting of infected cell lysates with antibodies to either Ser51-phosphorylated eIF2α or total eIF2α revealed no differences between WT and the three mutants (Supp. Fig. S4). We therefore concluded that eIF2α phosphorylation by PKR is not implicated in the downstream effects on HCV assembly. Activation of NFκB results in translocation of the p65 subunit from the cytoplasm to the nucleus. To test whether this downstream effector was responsible for the PKR effect on virus assembly we analysed infected cells by immunofluorescence with an antibody specific to p65 (Supp. Fig. S5). As expected, treatment with the NFκB activator TNFα resulted in efficient nuclear translocation of p65, however as shown in the representative images in Supp. Fig. S5 no such translocation was observed for either mock-infected or HCV-infected cells (WT or mutants). We thus conclude that activation of NFκB by PKR is not required for its effects on virus assembly, consistent with a requirement of PKR catalytic activity for the virus assembly block (Fig 7F).

We finally focused on IRF1 which drives expression of antiviral interferon-stimulated genes (ISGs) (60). IRF1 has been previously demonstrated to negatively regulate HCV genome replication (61), and the silencing of either IRF1 itself, or its effector targets PSMB9, APOL1 and MX1 enhanced HCV genome replication (62). The effects of IRF1 on HCV assembly have not been evaluated. To test this, we silenced IRF1 in Huh7.5 cells using CRISPR/Cas9 (Fig 14A), and electroporated these cells with WT or the mutant HCV RNAs. As was observed for PKR silenced cells, genome replication (Fig 14B) and viral protein production (Fig 14C, D) was unaffected by the lack of by IRF1 knockout. Reassuringly, when we analysed the assembly and release of the mutants, the production of infectious virus by C142A and E191A were restored to the same levels as WT and C190A in IRF1 knockout Huh7.5 cells (Fig 14E). These data confirmed that the ability of PKR to inhibit the assembly of HCV is mediated by its activation of the downstream effector IRF1.

**Fig 14.**
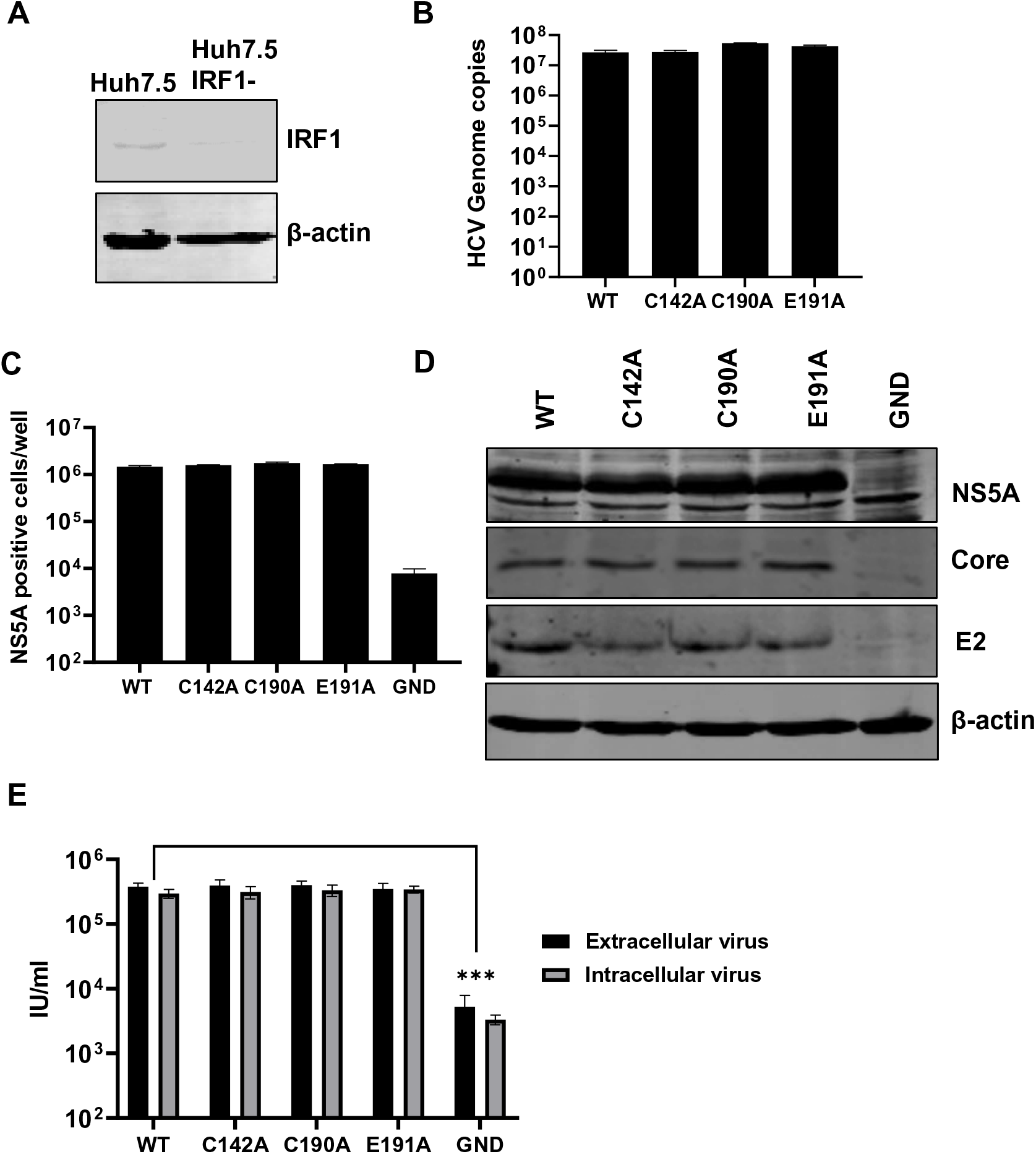
Virus assembly in IRF1-silenced Huh7.5 cells. **(A)** IRF1 expression was detected in silenced Huh7.5 cells by western blotting. Huh7.5 cells were electroporated with mJFH-1 WT and DI mutant C142A, C190A and E191A RNAs, together with an NS5B GND mutant as negative control. Virus genome replication was analysed detected directly by quantification of genome copies in cell lysates using qRT-PCR (**B**), and indirectly by enumerating NS5A positive cells at 72 hpe using the IncuCyte S3 (**C**). **(D)** Cell lysates were collected at 72 hpe and then analysed by western blot with the indicated antibodies. **(E)**. Extra- and intracellular virus harvested at 72 hpe were titrated onto Huh7.5 cells and quantified using the IncuCyte S3.

## Discussion

This study builds on our previously published work (1) and provides further evidence that DI of NS5A plays a key role in HCV assembly. However, our analysis revealed that only a limited number of residues are involved in this role as only two further residues (C142 and E191) share the previously identified function exhibited by V67 and P145. In particular we observed that although a significant patch of highly conserved residues is present on the surface of the NS5A DI monomer in proximity to P145, this is dispensable for virus assembly. A cluster of residues (P102, Y106, W111, and F149) surrounding P145 on three sides appear critical for genome replication. This is consistent with NS5A mediating a switch from virus replication to assembly, perhaps by interacting with a different subset of cellular and/or viral factors. The nature of the switch remains obscure although phosphorylation of the serine cluster in the low complexity sequence linking DI and DII has been proposed. As C142 and E191 are close to the C-terminus of DI it is conceivable that they are regulated by phosphorylation. It has been hypothesised that the two dimeric forms – open (1ZH1) and closed (3FQM) represent the two conformations of NS5A with different functions. In this regard it is interesting to speculate that in the closed dimer C142 is partially occluded within the dimer interface (Fig 1C) and more likely for the open form to function in virus assembly.

A role for DI in virus assembly is consistent with the observation that treatment of infected cells with an NS5A DAA (ledipasvir) inhibited virus assembly within 2 h. In contrast, inhibition of genome replication was not observed until >12 h, most likely due to the inability of NS5A DAAs to inhibit pre-existing replication complexes (17). Given that NS5A DAAs target DI, the implication of this data are that DI plays a role in assembly, as well as genome replication.

Our data show that for WT and C190A that efficiently assemble into infectious virus particles, the number of LDs per-cell is lower in comparison to mock infected cells (Fig 5). This was concomitant with an increase in the size of LDs, suggesting that virus infection coalesces LDs into larger entities that support the process of virus assembly. LDs were also distributed throughout the cytoplasm compared to the restricted perinuclear distribution observed in mock infected cells (Fig 6). These changes were not apparent for assembly defective mutants for which, the colocalisation of NS5A and Core with LDs, and the NS5A:Core colocalisation, were all reduced. Together, these data are consistent with the hypothesis that NS5A DI regulates the recruitment of both NS5A and Core to LDs, and modulates LD morphology and distribution to facilitate virus assembly.

Our observation that the silencing of PKR restored the phenotype of the two assembly defective mutants provides clues to the mechanism of action of DI in regulating assembly. NS5A binds viral RNA (21) and the open form (15) presents a basic surface in the groove between the monomers that is a possible RNA binding motif (Fig 1B). An attractive hypothesis is therefore that NS5A is involved in transporting nascent genomic RNA from sites of replication to sites of assembly (63, 64). In this scenario, the LD is a waystation on the route and at that point NS5A could deliver the RNA to the Core protein, rather like a baton in a relay race. One potential consequence of this is that the RNA would be transiently exposed in the cytosol, permitting detection by innate cytosolic sensors such as PKR. PKR is activated by binding to short (30 bp) dsRNA elements but can activated by imperfect dsRNA or single stranded RNA (65). In this regard, the HCV 5’ IRES has been shown to be both a potent activator (66) and inhibitor (67) of PKR. We postulate that DI interferes with the binding of PKR to nascent genomes, possibly by direct binding to PKR (Fig 13), preventing PKR activation and the induction of downstream antiviral pathways. PKR silencing also restored the LD phenotype (Figs 8 and 9), suggesting that activated PKR functions at the level of LD morphology to block virus assembly.

Our data suggest that neither eIF2α phosphorylation, nor activation of NFκB, are involved in the block to virus assembly (Supp. Figs S4 and S5), although these analyses were performed at 72 hpe and we cannot rule out transient effects at earlier times. PKR also activates IRF1, the silencing of which also restores the function of the virus assembly mutants. The implication of this observation is that one or more proteins whose expression is IRF1-dependent function to block virus assembly. In this regard a recent study (62) identified a number of IRF1 regulated genes (PSMB9, ApoL1 and MX1), which when silenced enhanced HCV replication approximately 3-fold. Overexpression of PSMB9 (a component of the proteosome) led to a 10-fold increase in HCV virus production, but a specific effect on virus assembly for ApoL1 and MX1 was not investigated. Whilst these factors may play a role, it is unlikely that they explain the 1000-fold reduction in virus production exhibited by C142A, which is completely restored by either PKR or IRF1 silencing. Of note ApoL1 (apolipoprotein L1) is an LD associated protein and, whilst other apolipoproteins have been implicated in HCV assembly (eg ApoE), the role of ApoL1 has not been investigated (68). The recruitment of ApoL1 to LDs is regulated by DGAT-1 (69), which is required for HCV assembly and recruits both NS5A and Core to LDs (29, 70). ApoL1 may therefore be an antiviral effector induced by PKR that acts on LD morphology to block virus assembly.

In conclusion, we here provide further support for a role of NS5A DI in controlling the assembly of infectious HCV particles. Analysis of assembly-defective DI mutants revealed a hitherto unknown antiviral pathway controlled by PKR and involving the downstream effector IRF1 which results in a block to virus assembly. Current work in our laboratory is focussed on dissecting the molecular mechanisms of this block to virus assembly and identifying the key players mediating these effects.

## Materials and Methods

### Cell lines

Huh7 (human hepatocellular carcinoma) (71) and Huh7.5 cells (a derivative of Huh7 from which a stable subgenomic replicon was ‘cured’ by IFN treatment and which exhibit a defect in RIG-I) (72) were used for electroporation. HEK-293T cells were used for transfections. Cells were cultured in Dulbecco’s Modified Eagles Medium (DMEM; Sigma) supplemented with 10 % fetal bovine serum (FBS), 100 IU penicillin/ml, 100 µg/ml streptomycin and 1 % non-essential amino acids (Lonza) in a humidified incubator at 37°C, 5 % CO_2_.

### Plasmid and virus constructs

Sub-genomic replicons with a luciferase reporter (mSGR-luc-JFH-1) and infectious virus (mJFH-1) were described previously (49). In both constructs unique BamHI/AfeI restriction sites flank the NS5A coding sequence. NS5A mutations were constructed using Q5 Site-Directed Mutagenesis Kit (New England BioLabs; E0554S) and cloned into either mSGR-luc-JFH-1 or mJFH-1 via the BamHI/AfeI restriction sites. NS5A domain I (amino acids 35 to 215) with mutations was PCR amplified and then cloned into pET-28a-Sumo vector using BamHI/XhoI restriction sites to construct pET-28a-Sumo-NS5A DI and mutants. Lentivirus constructs for silencing of PKR and IRF1 were obtained from Prof. Greg Towers (UCL) (54). PKR was amplified from Huh7 RNA and cloned into pcDNA3.1 vector using BamHI/XhoI restriction sites to construct pcDNA3.1-PKR. Primer sequences are available on request.

### Antibodies

Rabbit anti-Core (polyclonal serum R4210) was obtained from John McLauchlan (Center for Virus Research, Glasgow), mouse-anti E2 (AP33) was obtained from Arvind Patel (Center for Virus Research, Glasgow). The following antibodies were also used: sheep-anti NS5A (in house polyclonal antiserum) (73), mouse-anti β-Actin (Sigma Aldrich; A1978), rabbit anti-PKR (Abcam; ab32035), rabbit anti-phospho-PKR T446 (Abcam;ab32036) mouse anti-dsRNA J2 (Scicons; 10010200), mouse-anti His (BIO-RAD; MCA1396GA), mouse-anti p65 (F-6) (Santa Cruz; sc-8008), rabbit anti-eIF2α (Cell Signaling; 9722S), rabbit anti-phospho eIF2α (Cell Signaling; 9721S) and rabbit anti-IRF1 (Cell Signaling; 8478S). Secondary IRDye 680 and 800 labelled antibodies were obtained from LI-COR, AlexaFluor-conjugated 488, 594 and 647 antibodies and BODIPY (558/568)-C12 dye were obtained from ThermoFisher Scientific.

### Electroporation and luciferase assay

Huh7/Huh7.5 cells were washed in ice-cold PBS. Cells (5×10^6^) were resuspended in ice-cold PBS and electroporated with 2 µg of RNA at 950 µF, 270 V. Cells were resuspended in complete media and then seeded separately into 96-well plates at 3×10^4^ cells/well, or 6-well plates at 3×10^5^ cells/well. At 4, 24, 48 and 72 h post-electroporation (hpe), cells were harvested into 30 µl or 200 µl passive lysis buffer (PLB; Promega), incubated for 15 min at room temperature and stored at −80°C until used. Luciferase activity was assessed (Promega) on a FluoStar Optima luminometer. Data were recorded as relative light units (RLU).

### IncuCyte S3 analysis

Following immunofluorescence staining for NS5A, plates were detected using an IncuCyte S3 (Essen BioScience). Viral titres were obtained by analysing the total number of virus positive cells per-well for each dilution. As this method measures the absolute number of infected cells, rather than the number of foci of infected cells, the titres are represented as infectious units per mL (IU/mL).

### qRT-PCR

Total cellular RNA was harvested by lysis in TRIzol (Invitrogen) and extracted using chloroform. cDNA were synthesised from 1 µg RNA by reverse transcription using LunaScript RT SuperMix Kit (NEB; E3010). qRT-PCR was performed using Luna Universal qPCR Master Mix (NEB; M3003) with SYBR Green. Amplification was performed using the following primers: JFH-1 Forward: 5’- TCTGCGGAACCGGTGAGTA-3’ JFH-1 Reverse: 5’-TCAGGCAGTACCACAAGGC- 3’.

### Western blotting

Cells were washed twice in ice-cold PBS and cell lysates were harvested in 1 x GLB (1% Triton X-100, 120 mM KCl, 30 mM NaCl, 5 mM MgCl_2_, 10% glycerol (v/v), and 10 mM piperazine-N,N’-bis (2-ethanesulfonic acid) (PIPES)-NaOH, pH 7.2) with protease and phosphatase inhibitors (Roche; 5892791001). A total of 10 or 20 µg of each sample were denatured at 95°C for 5 min and separated by SDS-PAGE. Proteins were transferred to polyvinylidene fluoride (PVDF) membrane and blocked with 50% (v/v) Odyssey blocking buffer (LI-COR) diluted in 1X Tris-buffered saline with Tween-20 (TBS-T) (50 mM Tris-HCl pH 7.4, 150 mM NaCl, 0.1% Tween-20). Membranes were probed with primary antibodies (1:1000 dilution) at 4°C overnight and stained with IRDye labelled anti-mouse (700 nm) and anti-rabbit (800 nm) secondary antibodies for 1 h at room temperature (RT). Membranes were imaged on a LI-COR Odyssey Sa Imager.

### Titration of Virus

Huh7.5 cells were seeded into 96-well plates (3000 cells/well) and incubated at 37°C overnight. The following day, freeze-thawed intracellular virus was centrifuged at 16000 rpm for 5 min and the pellets were removed. Intracellular and extracellular virus samples were serially diluted two-fold and added to Huh7.5 cells. At 48 h cells were fixed in 4% (w/v) paraformaldehyde (PFA) for 30 min and washed twice with PBS. Cells were permeabilised with 0.25% Triton X-100 (Sigma-Aldrich) in PBS for 8 min and blocked with 3% BSA diluted in PBS. Cells were probed with primary antibodies (sheep anti-NS5A 1:2000) in PBS/3% BSA at 37°C for 1h. Cells were washed 5 times in PBS and incubated with Alexa Fluor-594 donkey anti-sheep (1:750) in PBS/3% BSA at RT for 1h. Plates were analysed using IncuCyte S3 software.

### Immunofluorescence analyses

Cells were seeded onto 16 mm glass coverslips in 12 well plates, fixed, permeabilised and blocked as described above. Cells were probed with sheep anti-NS5A (1:2000), rabbit anti-Core (1:500) or mouse anti-dsRNA J2 (1:200) in PBS/3% BSA at RT for 2 h. After washing three times, cells were incubated with Alexa Fluor-488 or -647 conjugated secondary antibodies (1:500 in PBS/3% BSA) at RT in the dark for 1 h. Lipid droplets were stained with BODIPY (558/568)-C12 dye (1:1000). Cells were mounted on to glass slides with Prolong Gold antifade reagent (Invitrogen) containing 4’,6’-diamidino-2-phenylindole dihydrochloride (DAPI) and sealed with nail varnish. Confocal images were acquired using a Zeiss LSM880 upright microscope with Airyscan. Post-acquisition analysis of images was performed using Fiji ImageJ (v1.49) software (74).

### Quantification of LD size and distribution

LD and DAPI channels were visualised using Fiji software and merged. LD size and distribution were analysed using the Analyse Particles module of Fiji. Data were analysed using GraphPad Prism and compared using two-tailed Student’s t tests.

### Co-localisation analysis

Overlap coefficients were analysed by Just-Another Co-localisation Plugin (JACop) in Fiji software from 10 cells derived from three independent experiments. Data were exported to GraphPad Prism and analysed using two-tailed Student’s t tests.

### Lentivirus production and construction of stable knockout cell lines

HEK293T cells in 10 cm dishes were transfected with 1 μg packaging plasmid p8.91, 1 μg envelope plasmid pMDG encoding VSV-G protein and 1.5 μg transfer plasmid lenti-CRISPRv2 encoding sgRNA to either PKR or IRF1 as described (54). Lentivirus supernatants were collected at 48 h and filtered through a 0.45 μm syringe. Huh7.5 cells were seeded into 6 well plates at density of 2.5 x 10^5^ cells/well and transduced with 1 ml/well lentivirus and 8 μg/ml polybrene for 24 h. Transduced cells were selected using 2.5 μg/ml puromycin at 72 h post transduction. Loss of target protein expression was confirmed by western blotting.

### Co-immunoprecipitation (Co-IP) assay

Cells were seeded into 6 well plates and lysed at 72 hpe using IP buffer (25 mM Tris-HCl pH 7.4, 150 mM NaCl, 1% NP-40, 1 mM EDTA; 5% glycerol) containing protease inhibitors for 1 h on ice. Lysates were centrifuged (13,000 × g for 5 min at 4°C) and supernatants are precleared with 10 μl protein G beads (Invitrogen; 1004D) at 4°C for 1 h. Cell lysates were then incubated with anti-PKR or anti-phospho-PKR antibodies at 4°C overnight prior to addition of 20 μl protein G beads for 2 h at 4°C. After extensive washing immunoprecipitated proteins were heat denatured at 95°C for 5 min, separated by SDS-PAGE and analysed by western blot.

### His-SUMO tagged NS5A expression, purification and pulldown assay

Plasmids were freshly transformed into *Escherichia coli* BL21 (DE3) pLysS and grown at 37 °C until OD_600_ values reached 0.6-0.8. Protein expression was induced by 100 μM isopropyl β-D-1-thiogalactopyranoside (IPTG) at 18°C for at least 6 h. Cells were recovered by centrifugation at 8000 x g for 15 min and resuspended in 50 ml binding buffer (100 mM Tris pH 8.2, 200 mM NaCl, 20 mM imidazole) supplemented with 40 μl DNase, 40 μl RNaseA, 2 mg/ml Lysozyme and protease inhibitors (Roche) per 1 L of pelleted culture. After incubation on ice for 30 min, samples were sonicated at an amplitude of 10 microns for 12 pulses of 20 sec separated by 20 sec on ice. After centrifugation at 4000 x g for 1 h at 4°C twice, supernatants were filtered through a 0.45 μm syringe filter. Samples were transferred to a 1 ml HisTrap FF column (Cytiva; 17531901) equilibrated with binding buffer and the column was washed 3 times using 5 column volumes of binding buffer. The samples were eluted with binding buffer containing 250 mM imidazole and dialyzed against 20 mM Tris-HCl, pH 8.2, 150 mM NaCl and 10% (v/v) glycerol.

His-SUMO tagged NS5A DI (5 μg) was diluted using 1x binding & wash buffer (BWB: 50 mM sodium phosphate pH 8.0, 300 mM NaCl, 0.02% Tween-20) and added to 20 μl Dynabead His-Tag beads (ThermoFisher Scientific; 10103D). After incubation on a roller at 4°C for 1 h, the beads were washed 5 times with BWB at 4°C. Lysates from pcDNA3.1-PKR transfected HEK293 cells were diluted with 1x pulldown buffer (3.25 mM Sodium-phosphate pH 7.4, 70 mM NaCl, 0.02% Tween-20) and added to the His-Tag beads at 4°C for 2 h. Beads were washed 5 times with BWB at 4°C, heat denatured at 95°C for 5 min, separated by SDS-PAGE and analysed by western blot.

### Statistical analysis

Statistical analysis was performed using an unpaired two-tailed Student’s t tests on GraphPad Prism version 9.30. **** (P<0.0001) and ** (P<0.01) indicate significant difference from wild type (n>=3). Data in histograms are displayed as the means ± S.E.

## Acknowledgments

We thank Prof. Greg Towers (University College, London) for the PKR, CypA and IRF1 silencing lentivirus constructs, Prof John McLauchlan (CVR, Glasgow) for the anti-Core antibody and Prof Arvind Patel (CVR, Glasgow) for the AP33 anti-E2 antibody. We thank Dr Niluka Goonawardane and Dr Ruth Hughes for help in Fiji ImageJ analysis.

## Supporting Materials

**Supp. Fig. S1 Virus assembly phenotypes in Huh7.5 cells.** Huh7.5 cells were electroporated with mJFH-1 WT and the DI mutant RNAs as indicated, together with an NS5B GND mutant as negative control. Extracellular virus harvested at 72 hpe was titrated in Huh7.5 cells and quantified using the IncuCyte S3.

**Supp.Fig S2. Location of mutated residues in DI.** The three surface exposed residues C142A, C190A and E191A proximal to P145 are displayed in two NS5A DI (genotype 1b) structures 1ZH1 (**B)** and 3FQM (**C).** Images on the right are zoomed into the boxed region shown in both space fill and ribbon format. Note that the disulphide bond formed by C142A and C190A was only observed in 1ZH1.

**Supp. Fig S3. Genome replication and virus assembly phenotypes in Huh7.5 silenced for CypA or PKR. (A)** In vitro transcripts of mSGR-luc-JFH-1 WT were electroporated into Huh7.5, Huh7.5 CypA or PKR silenced cell lines. Luciferase activity was measured at 72hpe. (**B)** In vitro transcripts of mJFH-1 WT were electroporated into Huh7.5, Huh7.5 CypA or PKR silenced cell lines. Extracellular infectious virus was titrated at 72 hpe.

**Supp. Fig S4. Expression of eIF2α and phospho-eIF2α.** Huh7.5 cells were electroporated with mJFH-1 WT and DI mutant C142A, C190A and E191A RNAs, together with an NS5B GND mutant as negative control. Cells were harvested at 72 hpe and lysed with GLB. eIF2α and phospho-eIF2α was analyzed by western blotting.

**Supp. Fig. S5 NF-κB activation in Huh7.5 cells infected with mJFH-1 or DI mutants.** Huh7.5 cells were electroporated with mJFH-1 WT and DI mutants C142A, C190A and E191A RNAs. At 72 hpe, cells were fixed and stained with mouse anti-P65 (green), sheep anti-NS5A (red) and DAPI. As a positive control to activate the NF-κB pathway uninfected Huh7.5 cells were treated with TNF-α for 24h. Mock: uninfected Huh7.5 cells.

**Supp. Fig S6. Immunofluorescent images of uninfected Huh7.5 cells. (A)** Representative image of mock infected cell from Fig 4 stained at 72 hpe with sheep anti-NS5A (white), rabbit anti-Core (green), BODIPY 558/568-C12 (red) and DAPI. Scale bar 5 μm. **(B)** Representative image of mock infected cell from Fig 8 stained at 72 hpe with sheep anti-NS5A (white), rabbit anti-Core (green), BODIPY 558/568-C12 (red) and DAPI. Scale bar 5 μm. **(C)** Representative image of mock infected cell from Fig 11 stained at 72 hpe with sheep anti-NS5A (green), mouse anti-dsRNA J2 (white), BODIPY 558/568-C12 (red) and DAPI. Scale bar 5 μm.

